# Seroprevalence of IgG Antibody against SARS-CoV-2 Nucleocapsid protein and Associated Risk Factors

**DOI:** 10.1101/2023.02.09.527802

**Authors:** Yeamin Farabi Chowdhury, Faruk Hossen, S. M. Rashadul Islam, Md. Saddam Hossain, Kazi Mahtab-Ul-Islam, Sumaiya Islam Chowdhury, Md. Rakibul Hasan, Nishat Tasnim, Sharmin Sultana, Md. Aftab Ali Shaikh, Md. Rezaul Karim

## Abstract

Estimation of antibody development against SARS-CoV-2 is essential means for understanding the immune response against the virus. We reported IgG antibody development status against Nucleocapsid protein of the virus and compared with lifestyle (health and food habits), co-existing diseases, vaccination and COVID-19 infection status. ELISA (Enzyme Linked Immunosorbent Assay) was performed to assess IgG antibodies targeted against the Nucleocapsid protein of SARS-CoV-2 in participants (n=500). In this seroprevalence study, serological data were estimated for a period of 10 months in the participants who were aged 10 years and above. Sociodemographic and risk factors related data were collected through a written questionnaire and chi-square test was performed to determine the association with seropositivity. The overall seroprevalence of anti-SARS-CoV-2 antibodies among the study subjects was 47.8%. Estimates were highest among the participants of 21-40 years old (55.1%), and lowest in older aged (>60 years) participants (39.5%). Among the Sinopharm vaccinated individuals 81.8% had developed anti-Nucleocapsid antibody. Physical exercise and existence of comorbidities like hypertension and diabetes were the distinguishing factors between seropositive and seronegative individuals. Seropositivity rate largely varied among symptomatic (67%) and asymptomatic (33.1%) COVID-19 infected participants. The findings suggest that residents of Dhaka city had a higher prevalence of anti-nucleocapsid antibody in the second year of the pandemic. This indicates the improvement of immunological status among the population. Finally, the study emphasizes on maintaining active and healthy lifestyle to improve immunity. However, the absence of IgG antibodies in many cases of COVID-19 infected individuals suggests that antibodies wane with time.

**Key messages:**

- The overall seroprevalence of anti-Nucleocapsid IgG among the study subjects was determined to be 47.8%.
- Age, regular physical exercise, existence of comorbidities were the identified parameters associated with seroprevalence.
- This study observed lower prevalence of Anti-Nucleocapsid antibody among asymptomatic cases of COVID-19 infected individuals compared to symptomatic cases.

## Introduction

The extremely infectious Coronavirus Disease-2019 (COVID-19) was originally discovered in Wuhan, Hubei Province, China, caused by the new RNA virus SARS-CoV-2 [1]. Within three months of the outbreak, the virus had spreaded all over the world and was declared a pandemic by the World Health Organization (WHO) in March 2020. Bangladesh, with its inadequate medical facilities and dense population, remains one of the most vulnerable countries throughout the epidemic [2]. COVID-19 was initially confirmed in Bangladesh on March 8, 2020. Until 27 March 2022, a total number of 1.95 million people have been infected and 96.67 million people have received two doses of vaccine [3].

Real Time Polymerase Chain Reaction (RT-PCR) diagnoses viral RNA in symptomatic patients to monitor SARS-CoV-2 infection in individuals. Cases that are mildly symptomatic or asymptomatic often are excluded from reports. As a result, true SARS-CoV-2 infection rates differ from reported cases [4]. Serological tests are necessary to identify all infected persons, regardless of clinical symptoms to undertake surveillance and limit the viral spread [5].

When infected with SARS-COV-2, the virus triggers a humoral immune response that results in the production of antibodies against Spike glycoprotein (S) and Nucleocapsid protein (N) within the first 7 days after onset of symptoms [5–9]. IgG and IgM antibody development in SARS-CoV-2 infection occurs concurrently or sequentially which differs from classical infection dynamics where IgM is expected to appear first and IgG is detected in delayed time [6]. Long-term immunity, however, necessitates the formation of IgG [10]. The speed of viral clearance is determined by the dynamics of the humoral immune response. Rapid viral clearance is linked to earlier antibody responses since low initial SARS-CoV-2 RNA was found in individuals who lack serum IgG. This report by Masiá et al., (2021) suggested that the intensity of viral replication influences the production of adaptive humoral responses. The distribution and variance of antibody dynamics are considered to be linked to age, gender, co-existing disease, viral load, and other parameters that influence disease severity [12].

Antibody responses to pathogens in serum or plasma are measured using serological tests. Different methods, including as ELISA, lateral flow assays, and Western Blot assays, are used to perform serological tests. These assays are useful for determining protective antibody titers in the event of infection or immunization [13]. Reinfection or illness severity is linked to decreasing antibody levels against SARS-CoV-2 [14]. It is important to determine the underlying cause behind seronegative individuals with a history of infection. Over a 10-month period, we investigated antibody production against the Nucleocapsid protein of SARS-CoV-2 in 500 people and correlated seropositivity with numerous risk factors. Antibodies against the nucleocapsid protein are mostly observed in persons after they had been infected with the virus and have been vaccinated with vaccines that use the virus in its whole attenuated form [15]. We surveyed the individuals on their lifestyles, food habits, COVID-19 infection status and comorbidities, and then compared them to their IgG antibody development.

## Methods

### Study Subjects

The research was carried out at the Institute of Technology Transfer and Innovation, Bangladesh Council of Scientific and Industrial Research in Dhanmondi, Dhaka. A total of 500 study subjects participated in the study who was mostly residents of Dhaka city. A standardized questionnaire was given to each participant which asked about socio-demographic characteristics, lifestyle, dietary and healthy habits, existing co-morbidities, COVID-19 infection status and vaccination status. Before registering for the survey, the participants had to provide verbal and written consent.

### Sample collection

The experiment was performed using fresh serum. Freshly collected peripheral venous blood of 3 ml was drawn from each participant and kept in a clot activator blood collection tube. After 5-10 minutes, clot formed in the blood and the tubes were centrifuged at 2,000 rpm for 8 minutes. Serum was collected in Eppendorf from the upper aqueous portion of centrifuged blood.

### Laboratory analysis

Anti-SARS-CoV-2-N-IgG was measured in the lab using a commercially available ELISA kit named ID Screen SARS-CoV-2-N IgG Indirect by Innovative Diagnostics that targets the virus’s Nucleocapsid protein. According to the test protocol, 1:20 diluted serum was used in the experiment. The ELISA plate from the commercial kit was coated with Nucleocapsid antigen of the virus to which only specific antibody present in the serum bound. The final optical density was determined by Multiskan ELISA plate reader.

### Statistical analysis

SPSS software version 21 was used for all statistical analyses. The Chi-square test was performed to examine the association between seropositivity and risk factors (age, sex, profession and lifestyle). Variables with a p-value of less than 0.05 were considered statistically significant.

## Results

Among the participants, 158 (68.4%) were male and rest were female (31.6%). 11 (2.2%) of the participants were of age 10 to 20 years, 225 (45%) were of age 21-40 years, 229 (45.8%) were of 41-60 years and 35 (7%) were above 60 years old. Among the participants, 142 (28.4%) were health professionals, 20 (4%) were engineers, 19 (3.8%) were teachers, 26 (5.2%) were businessmen, 73 (14.6%) were scientists and rest 220 (44%) were of other professions that included government and private service holders, bankers, homemakers, students etc.

### Demographics of the study participants

The socio-demographic characteristics of the participants are presented in Table 1. This data represents that the study subjects were mainly working-class people who had to go outside the house during pandemic. The highest prevalence of antibody (55.1%) was among the age group of 21-40 years. Out of the 43 individuals in the age group above 60 years, 17 individuals had developed antibody which was found the lowest prevalence (39.5%) of antibody among all the age groups. Seroprevalence of current smokers (35.16%) was significantly lower than that of never smokers (51.92%) (*P*=<0.05). The people who exposed to the dust had found highest prevalence (57.32%) of antibody than the people who did not expose to the dust (35.17%).

**Table 1:**
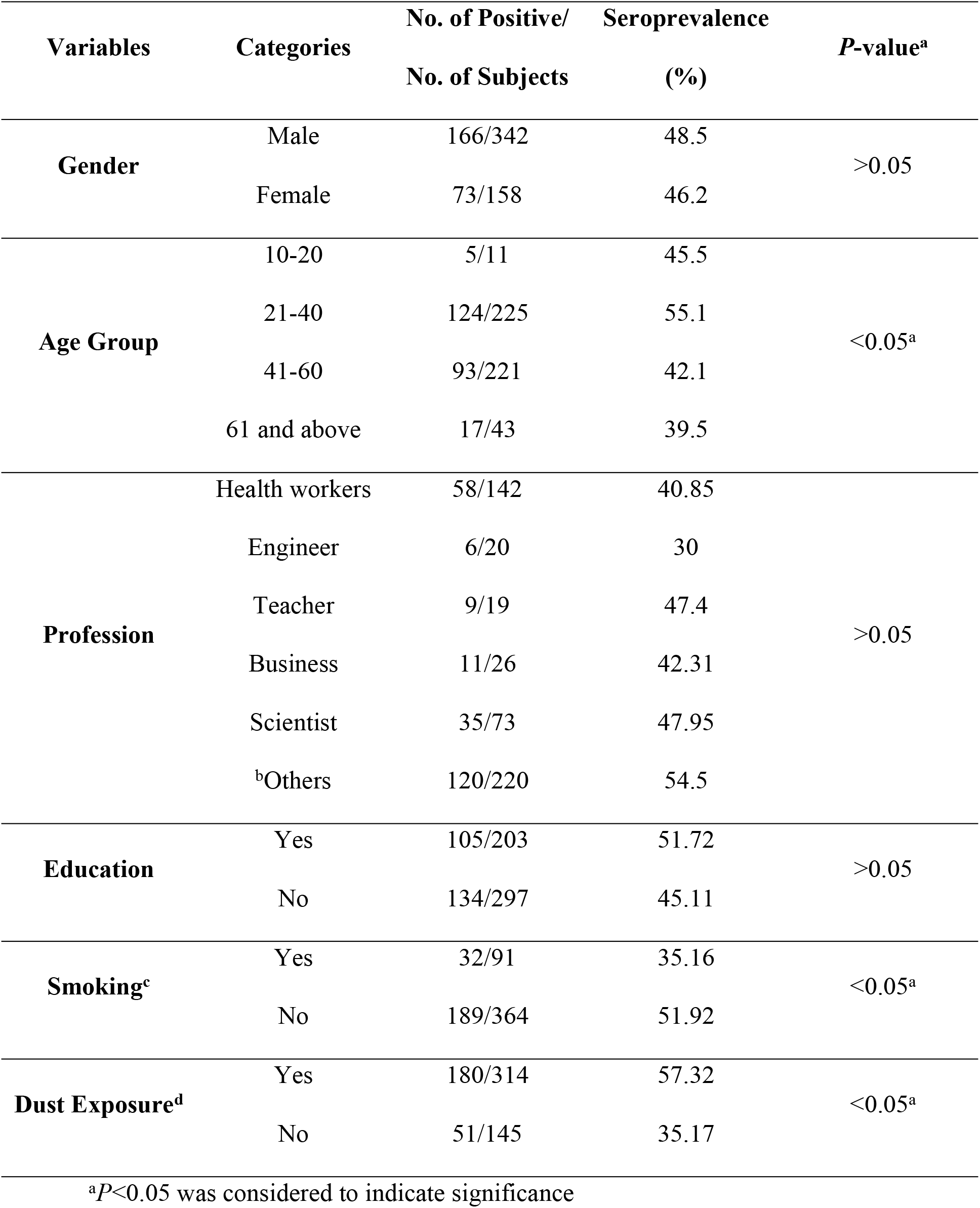
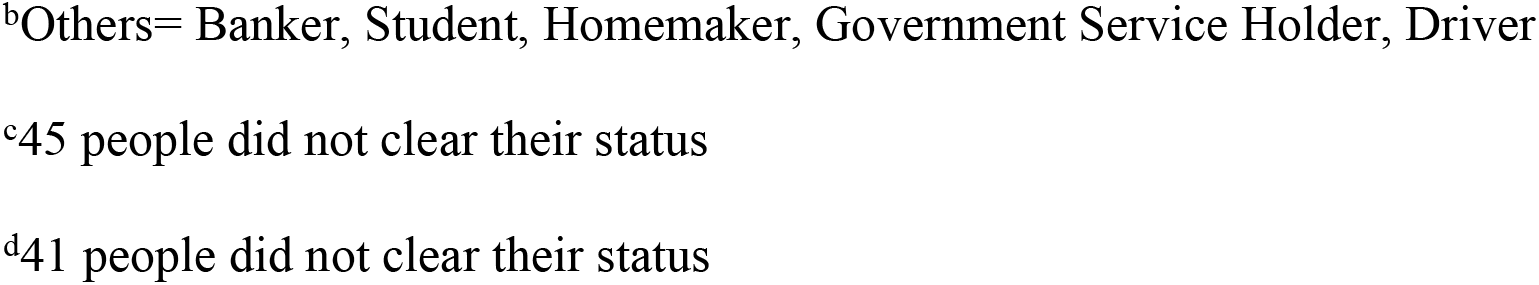
Seroprevalence of IgG antibody according to Socio-demographic characteristics of the study subjects

| Variables | Categories | No. of Positive/<br>No. of Subjects | Seroprevalence<br>(%) | *P*-value<sup>a</sup> |
| --- | --- | --- | --- | --- |
| **Gender** | Male | 166/342 | 48.5 | >0.05 |
|  | Female | 73/158 | 46.2 |  |
| **Age Group** | 10-20 | 5/11 | 45.5 | <0.05<sup>a</sup> |
|  | 21-40 | 124/225 | 55.1 |  |
|  | 41-60 | 93/221 | 42.1 |  |
|  | 61 and above | 17/43 | 39.5 |  |
| **Profession** | Health workers | 58/142 | 40.85 | >0.05 |
|  | Engineer | 6/20 | 30 |  |
|  | Teacher | 9/19 | 47.4 |  |
|  | Business | 11/26 | 42.31 |  |
|  | Scientist | 35/73 | 47.95 |  |
|  | <sup>b</sup>Others | 120/220 | 54.5 |  |
| **Education** | Yes | 105/203 | 51.72 | >0.05 |
|  | No | 134/297 | 45.11 |  |
| **Smoking<sup>c</sup>** | Yes | 32/91 | 35.16 | <0.05<sup>a</sup> |
|  | No | 189/364 | 51.92 |  |
| **Dust Exposure<sup>d</sup>** | Yes | 180/314 | 57.32 | <0.05<sup>a</sup> |
|  | No | 51/145 | 35.17 |  |

<sup>a</sup>*P*<0.05 was considered to indicate significance
<sup>b</sup>Others= Banker, Student, Homemaker, Government Service Holder, Driver
<sup>c</sup>45 people did not clear their status
<sup>d</sup>41 people did not clear their status

### Lifestyle and dietary habits with antibody positivity

In this study, the healthy habits and risk factors of participants were taken into consideration that might be responsible for the persistence of antibody after infection or vaccination. In Table 2, different habits and risk factors of subjects such as regular walking, regular exercise, dietary habits of consumption of vegetables, fruits, fast foods and vitamin-C had been demonstrated. Significantly higher (*P*<0.05) seropositivity of 52.1% and 54.4% was found among the participants who had the habit of regular walking and exercising respectively. The dietary habits were not found significantly (*P*>0.05) correlated with the antibody development.

**Table 2:** Association of IgG antibody with lifestyle, dietary and healthy habits of the study subjects

| Risk factors | Categories | No. of Positive/ No. of Subjects | Seroprevalence (%) | P-value<sup>a</sup> |
| --- | --- | --- | --- | --- |
| Regular Walking | Yes | 171/328 | 52.1 | <0.05<sup>e</sup> |
|  | No | 68/172 | 39.5 |  |
| Regular Exercise | Yes | 93/171 | 54.4 | <0.05<sup>e</sup> |
|  | No | 146/329 | 44.4 |  |
| Regular Vegetable Intake | Yes | 223/466 | 47.9 | >0.05 |
|  | No | 16/34 | 47.1 |  |
| Regular Fruits Intake | Yes | 195/393 | 49.6 | >0.05 |
|  | No | 44/107 | 41.1 |  |
| Regular Fast Food Intake | Yes | 40/85 | 47.1 | >0.05 |
|  | No | 199/415 | 48 |  |
| Vitamin C Intake | Yes | 193/395 | 48.9 | >0.05 |
|  | No | 46/105 | 43.8 |  |

<sup>a</sup> $P < 0.05$ was considered to indicate significance

### COVID-19 infection and vaccination status with antibody development

The Covid-19 infection and vaccination status of the participants has been demonstrated in Table 3. The study showed that the prevalence of antibody among 162 individuals who tested positive for Covid-19 was 67.9%. Only 54 (37.5%) individuals were found seropositive among 144 individuals who tested negative for Covid-19. Among the undiagnosed individuals 38.7% were found seropositive.

**Table 3:** Seroprevalence according to COVID-19 infection and Vaccination status

| Variables | Categories | No. of Positive/ No. of Subjects | Seroprevalance (%) | P-value<sup>a</sup> |
| --- | --- | --- | --- | --- |
| **COVID-19 Test Status** | Positive | 110/162 | 67.9 | <0.05<sup>a</sup> |
|  | Negative | 54/144 | 37.5 |  |
|  | Undiagnosed | 75/119 | 38.7 |  |
| **Vaccination Status** | Partially Vaccinated | 35/66 | 53.03 | >0.05 |
|  | Fully Vaccinated | 149/320 | 46.56 |  |
|  | Non Vaccinated | 55/114 | 48.25 |  |
| **Name of Vaccine** | AstraZeneca | 100/273 | 36.6 | <0.05<sup>a</sup> |
|  | Pfizer | 7/13 | 68.18 |  |
|  | Sinopharm | 54/86 | 81.8 |  |
|  | Moderna | 15/25 | 60 |  |
| **Coexisting disease** | Yes | 94/222 | 42.3 | <0.05<sup>a</sup> |
|  | No | 145/278 | 52.2 |  |
| **Coexisting disease (Specific)** | Diabetes | 35/94 | 37.2 | <0.05<sup>a</sup> |
|  | Hypertension | 38/113 | 33.6 |  |
|  | Other Diseases<sup>b</sup> | 33/68 | 48.5 |  |

<sup>a</sup> $P<0.05$ was considered to indicate significance
<sup>b</sup> COPD, liver disease, heart disease, renal disease, asthma, obesity, malignancy

The vaccination status of the participants showed that the subjects who received only 1^st^ dose of vaccine had higher seroprevalence of antibody (53.03%) against Nucleocapsid protein of COVID-19 compared to that of fully vaccinated (46.56%) individuals and non-vaccinated (48.25%) individuals.

In this study, people who received Sinopharm vaccine were found to develop significantly higher (*P*<0.05) level (81.8%) of antibody whereas significantly lower (*P*<0.05) level of antibody (36.6%) was found among the individuals vaccinated with AstraZeneca.

Significantly higher seropositivity (52.2%) was found among the individuals who did not have history of co-existing diseases like hypertension, diabetes, COPD, liver disease, heart disease, asthma and other diseases. On the other hand, significantly lower seropositivity (42.3%) was observed among the patients who had either one or more co-existing diseases. Significantly lower (*P*<0.05) antibody development was detected in the participants afflicted by Hypertension (33.6%) and Diabetes (37.2%).

### Association of seroprevalence with appearance of symptoms on COVID-19 infection

Seroprevelance in symptomatic and asymptomatic cases had been associated with COVID-19 test status of participants which is demonstrated in Table 4. The patients who had the symptoms of COVID-19 and tested positive for the virus had showed significantly (*P*<0.05) higher level (67%) of antibody in comparison with the negative (16.5%) and undiagnosed (16.5%) COVID-19 cases. Seropositivity was also significantly higher (40.5%) among the participants who neither tested nor experienced the conventional symptoms of COVID-19. Among the asymptomatic participants who tested positive for SARS-CoV-2 had developed antibody at a rate of 33.1% and who tested negative for SARS-CoV-2 had developed antibody at a rate of 26.4%. Antibody was not detected in a good number of participants (40%) who had experienced symptoms while also being diagnosed positive for COVID-19. Among the individuals who neither tested nor experienced the conventional symptoms of COVID-19 were found seronegative at a rate of 55.1% which was significantly higher (*P*<0.05) than the other asymptomatic seronegative outcomes.

**Table 4:** Seroprevelance in symptomatic and asymptomatic cases

| Seroprevalence | COVID-19 Symptom Appearance | COVID-19 Test Status |  |  | Total<br>(n=500) | P-value<sup>a</sup> |
| --- | --- | --- | --- | --- | --- | --- |
|  |  | Positive | Negative | Undiagnosed |  |  |
| Positive | Symptomatic | 61 (67%) | 15 (16.5%) | 15 (16.5%) | 91 | <0.05<sup>e</sup> |
|  | Asymptomatic | 49 (33.1%) | 39 (26.4%) | 60 (40.5%) | 148 |  |
| Negative | Symptomatic | 26 (40%) | 28 (43%) | 11 (16.9%) | 65 | <0.05<sup>e</sup> |
|  | Asymptomatic | 26 (13.3%) | 62 (31.6%) | 108 (55.1%) | 196 |  |

<sup>a</sup> $P<0.05$ was considered to indicate significance

### Prevalence of SARS-CoV-2 antibodies

The monthly overview of the antibody test results is presented in Fig 1. Over the course of the study, 47.8% participants were found to have antibodies against Nucleocapsid protein of the virus. Number of seropositive individuals significantly increased in the last 2 months. Antibody prevalence among the subjects in the 9^th^ and 10^th^ month were 64.2% and 75% whereas in the 3^rd^ month of the study antibody development was found in only 30.8% of the subjects.

**Fig 1:**
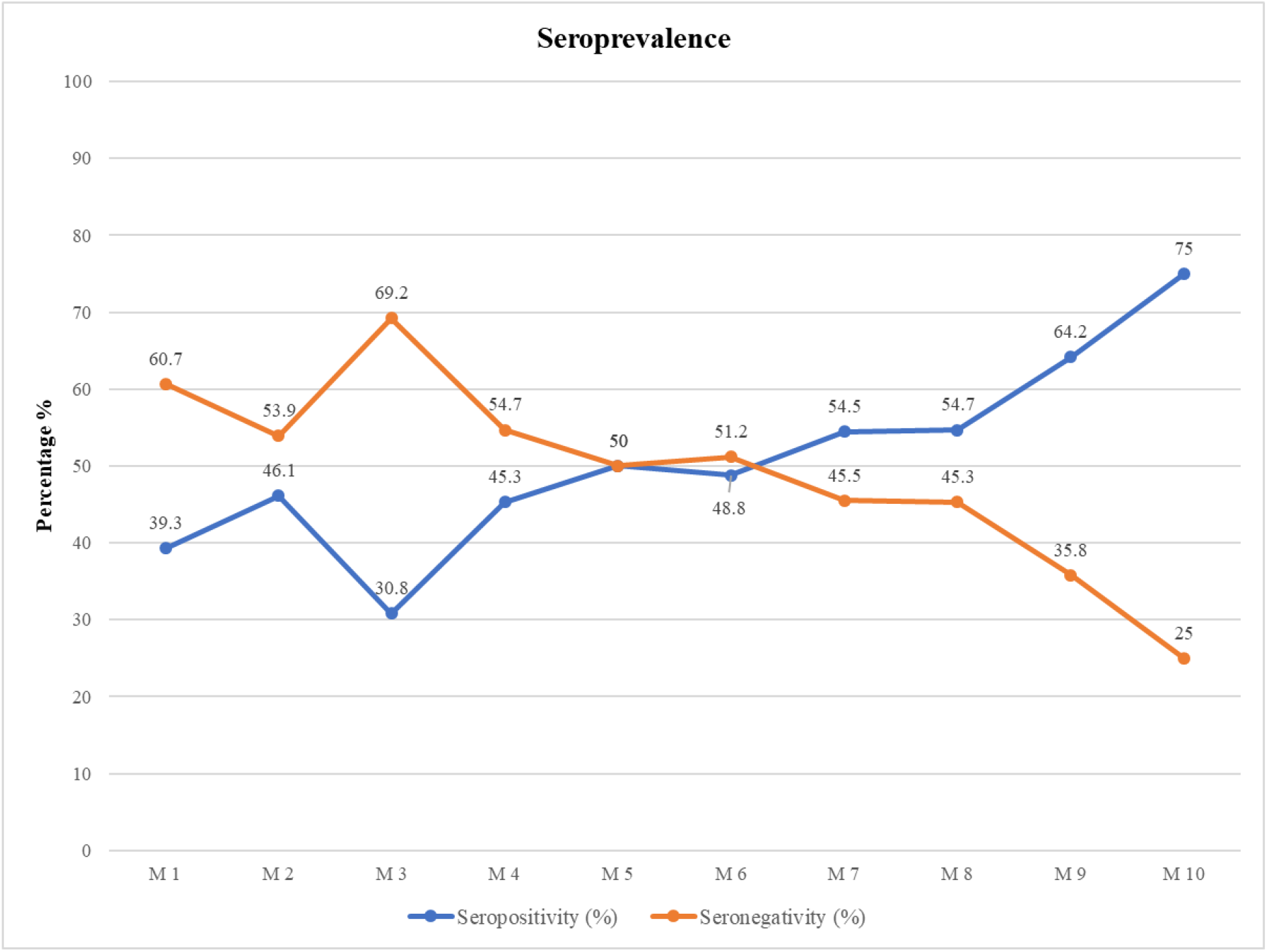
Seroprevalence estimates per month. Percentage of seropositivity is presented with blue line and that of seronegativity is presented in red line

## Discussion

As the presence of IgG antibody is associated with protection against the virus, the preliminary purpose of this study was to assess the status of COVID-19 antibody development among the residents of Dhaka city. The seroprevalence study was conducted while Bangladesh was experiencing the second wave of the pandemic. The present study was set to see a link between risk factors and IgG antibody formation and persistence of the produced antibody in the body [16].

There was no significant difference between male and female regarding IgG development. Participants aged >60 years had lower antibody level than patients age spanning 10-60 years (*P*<0.05). The highest seroprevalence found among the age group 21-40. This data suggests that older people have a lower response to developing protective antibodies [16] than younger individuals in significant quantities [17] and therefore, older people are at higher risk of developing severe symptoms [18]. This data is consistent with other seroprevalence surveys done during pandemics on different populations [19]. Compared to non-smokers, current smokers were found to have a lower seroprevalence (51.92% in non-smokers, in contrast to 35.16% seroprevalence in current smokers). Certain studies have indicated that smoking cigarettes impairs the immune system [20] and reduces production of IgA, IgG and IgM antibodies [21]. Even after vaccination, current smokers have shown significant lower antibody titre [22–26].

Regular walking and regular exercise which are actually considered as physical exercise were shown to have a substantial correlation with the seropositivity of the persons among the risk factors [27]. Aman & Masood, (2020) reported that a balanced and nutritious diet can help to build a strong immune system that can withstand the viral attack. People who eat a well-balanced diet appear to be healthier, with stronger immune systems and a lower risk of infections. However, healthy dietary habits like vegetable and fruits intake on regular basis were not found to have direct impact on IgG response of the subjects.

The efficacy of Sinopharm vaccine which uses whole attenuated virus [29] in its dosage were estimated in our study. The seropositivity of Sinopharm vaccinated subjects were found 81.8%. The individuals vaccinated by Pfizer, Moderna and Astrazeneca were found to have a moderate development of antibody in their body. These vaccines use mRNA of spike protein in their dosage. As we assessed antibody against Nucleocapsid protein of the virus, antibody development against mRNA vaccines were not found in satisfactory level. Among the mRNA vaccinated individuals, some have developed IgG antibody against Nucleocapsid protein through direct infection of SARS-CoV-2 either before or after vaccination [30]. No significant difference was found among the vaccinated and non-vaccinated subjects.

It has already been established that older age and certain comorbidities are risk factors for hospitalization and death by COVID-19 infection [31]. This study has showed that the existence of comorbidities lowers IgG response in the individuals which is supported by Hoque et al., (2021) who stated that the existence of comorbidity lowers antibody response in human body. This indicated that the older individuals over 60 years and individuals with co-morbidities are more likely to develop a defective immune response and suffer higher rates of morbidity and mortality [32]. Watanabe et al., (2022) stated that Hypertension were strongly associated with lower antibody titers which supports our findings that IgG developments were significantly low in patients suffering from hypertension (33.6%) and Diabetes (37.2%). In another study it had been found that some comorbidities, including hypertension, COPD, immunosuppression, and type 2 diabetes, were negatively linked with vaccine efficacy [34]. The seropositivity rate was higher in symptomatic than in asymptomatic COVID-19 infection (67% versus 33.1%; *P*<0.05). This observation is supported by previous studies where COVID-19 associated symptoms had higher IgG humoral response and antibody titers compared to asymptomatic patients [35–37]. Varying levels of antibody against anti-SARS-CoV-2 is linked to the severity of the disease [36, 38]. In symptomatic cases, IgG and IgM positively elevates and overall immune response develops shortly after infection [29, 39]. Interestingly, among the asymptomatic individuals seropositivity was higher in the undiagnosed individuals (40.5%) than the COVID-19 tested positive (33.1%) and negative (26.4%) individuals. This observation is supported by Shirin et al., (2020) who found lower antibody levels in people with asymptomatic infection. On the other hand, positive immune response was found in asymptomatic uninfected individuals either through vaccination [17] or immunity developed through previous infection with COVID-19 [15, 41]. In this study asymptomatic undiagnosed individuals showed higher immune response either they were silently infected but did not have any confirmatory test of COVID-19 or vaccination increased their immune status. Higher seronegativity (40%) was found in COVID-19 infected symptomatic individuals than the asymptomatic individuals (13.3%). Alongside, in undiagnosed symptomatic and asymptomatic participants, the seronegativity rate was 16.9% and 55.1% respectively. This can be caused by either impaired IgG response because of immune suppressive factors (such as Comorbidities, old age etc.) [17] or antibody reduction after a significant time. In the majority of people, IgG responses against SARS-CoV-2 appear to persist longer than 6 months [42–44] which is associated with age and disease severity [45]. A rise in seroprevalence was evident over the study’s ten months, with the peak occurring in the last month. Seropositivity went from 39.3% in the first month to 75% by the end of last month.

The serological study found that the prevalence of antibodies against Nucleocapsid protein of SARS-CoV-2 was found in 47.8% of the study subjects. This study shows that a large population had developed COVID-19 specific antibodies during the 2nd year of the pandemic while the nationwide vaccination program was going in full swing. Therefore, the data suggests that a vast majority of the population is protected against COVID-19 as long as antibody persists in the body.

There are some major limitations to this study. Firstly, we assessed the development of IgG against Nucleocapsid protein using the ELISA technique. This excludes the assessment of subjects who developed antibody through vaccination by vaccines that uses mRNA of Spike protein. As a result, the study excludes a substantial percentage of seropositive participants. Secondly, there was an over representation of working-class people of the age group 21-60. The study population wasn’t completely random rather the research concentrated on working class people who had gone out during the pandemic. Thirdly, the accuracy and credibility of serology assays are questioned because of the risk of false-positive instances due to cross-reaction with other coronavirus such as the common cold. Again, false negative results are also plausible. However, serology tests by ELISA yielded high specificity of around 99% [46]. Finally, this research also excluded the group of physically working people like rickshaw puller and daily laborer.

## Conclusion

Seroprevalence data are crucial for determining the pandemic’s scope and spread, as well as predicting the likelihood of subsequent waves. It addresses public health issues, such as the need for future lockdowns and the success of immunization campaigns, booster dosage requirements and the time interval between booster doses. However, the seroprevalence data also represent the efficiency of different vaccines, vaccinated among the individuals of our country.

## Data Availability

Data available on request. The data underlying this article will be shared on reasonable request to the corresponding author.

## Ethical approval

The study was approved by the Ethics Review Committee of Bangladesh Council of Scientific and Industrial Research (BCSIR), ref. no. 39.02.0000.011.37.017.2017/351, date: 15.01.2021.

## Acknowledgments

The authors are grateful to the Bangladesh Council of Scientific and Industrial Research (BCSIR) and the Ministry of Science and Technology, Bangladesh.

## Author contributions

FH: Conceptualization, Writing-reviewing and editing. YFC and SMRI: Methodology, Formal analysis, Investigation, Data curation, Writing-original draft preparation, Writing-reviewing and editing. SIC, MSH, KMUI and NT: Investigation, Writing-reviewing and editing. MRH: Methodology, Investigation, Writing-reviewing and editing. MRK: Investigation, Writing-reviewing and editing, Supervision. MAAS: Resources, Supervision. All authors read and approved the final manuscript.

